# A novel long noncoding RNA LINC00844 regulates prostate cancer cell migration and invasion through androgen receptor signaling

**DOI:** 10.1101/244459

**Authors:** Shreyas Lingadahalli, Sudhir Jadhao, Ying Ying Sung, Mi Chen, Lingling Hu, Xin Chen, Edwin Cheung

## Abstract

The majority of the human genome is transcribed, yielding a rich repository of non-coding transcripts that are involved in a myriad of biological processes including cancer. However, how non-coding transcripts such as long non-coding RNAs (lncRNAs) function in prostate cancer is still unclear. In this study, we have identified a novel set of clinically relevant androgen-regulated lncRNAs in prostate cancer. Among this group, we showed LINC00844 is a direct androgen-regulated target that is actively transcribed in androgen receptor (AR)-dependent prostate cancer cells. The expression of LINC00844 is higher in normal prostate compared to malignant and metastatic prostate cancer samples and patients with low expression demonstrate poor prognosis and significantly increased biochemical recurrence, suggesting LINC00844 may function in suppressing tumor progression and metastasis. Indeed, *in-vitro* loss-of-function studies revealed that LINC00844 prevents prostate cancer cell migration and invasion. Moreover, findings from gene expression analysis indicated that LINC00844 functions in *trans*, affecting global androgen-regulated gene transcription. Mechanistically, we provide evidence to show LINC00844 is important in facilitating AR binding to the chromatin. Finally, we demonstrated LINC00844 mediates its phenotypic effects in part by activating the expression of NDRG1, a crucial cancer metastasis suppressor. Collectively, our findings suggest LINC00844 is a novel coregulator of AR that plays a central role in the androgen transcriptional network and the development and progression of prostate cancer.

## Introduction

Prostate cancer is the most common non-cutaneous cancer affecting men worldwide [1]. Androgen receptor (AR) is a hormone-regulated transcription factor (TF) that occupies a central position in both normal prostate development and its carcinogenesis [2]. AR mediates its biological effect by controlling the transcription of downstream target genes [3, 4]. Therefore, anti-androgens which inhibit the transcriptional activity of AR have become a part of standard care in prostate cancer therapy. Regardless of the favorable initial response, a majority of the patients will relapse into a more lethal and incurable form called metastatic castration resistance prostate cancer (mCRPC) [5–7]. Extensive research over the years has established that the re-activation or aberrant activation of the AR signaling pathway remains a critical event in the development of mCRPC [8, 9]. Hence, it is imperative to elucidate the underlying molecular mechanisms of AR-mediated transcriptional regulation in order to develop better diagnostic markers and improved therapies for the management of advanced prostate cancer.

The discovery that the majority of the human genome is transcribed yielding a rich repository of non-coding RNAs (ncRNAs) has generated widespread interest in understanding their functional roles [10]. NcRNAs that are greater than 200 bp in length are arbitrarily classified as long non-coding RNA (lncRNA) and have been implicated in a myriad of biological processes including cancer [11, 12]. With respect to prostate cancer, a number of lncRNAs have emerged as prostate cancer biomarkers such as Prostate Cancer Antigen 3 (PCA3) [12, 13] and SChLAP1 [14]. Several recent reports have now implicated a critical role for lncRNAs in directly regulating the transcriptional activity of AR in prostate cancer. For example, Rosenfeld and colleagues showed two lncRNAs, Prostate Cancer Gene Expression Marker-1 (PCGEM1) and Prostate Cancer Noncoding RNA1 (PRNCR1), which are important for enhancing androgen-dependent transcription by promoting the chromatin looping of AR-bound enhancers to their target gene promoters [15]. In addition, CTBP1-AS, an AR-stimulated antisense lncRNA, was found to repress the expression of CTBP1 in a *cis-*regulatory manner as well as tumor-suppressive genes in *trans* by recruiting the histone deacetylase corepressor complex via the PTB-associated splicing factor (PSF) [16]. Similarly, HOTAIR was reported to physically interact with AR in order to prevent the E3 ubiquitin ligase-mediated degradation of the receptor and thus facilitating an androgen-independent AR transcriptional program and the promotion of mCRPC transformation [17]. Although we are beginning to understand the role of a few lncRNAs in prostate cancer, our current knowledge is still limited and a vast repository of lncRNAs have yet to be identified or extensively investigated with respect to AR-mediated transcriptional regulation and prostate cancer biology.

In this study, we have used a combination of bioinformatic analysis and molecular approaches to identify and functionally characterize a novel AR-regulated lncRNA in prostate cancer called Long Intergenic Long Non-Coding RNA 844 (LINC00844). We demonstrate that LINC00844 is clinically significant and potentially plays an important role in metastatic transformation of prostate cancer and provide evidence to show that LINC00844 inhibits migration and invasion of prostate cancer cells without altering the rate of cell proliferation. In addition, we establish that LINC00844 is a critical component of the AR transcriptional network and enhances the global expression of androgen-regulated genes by promoting AR recruitment to chromatin. Finally, our study also reveal that LINC00844 mediates its phenotypic effects in part by regulating the DHT-mediated activation of NDRG1, a known metastatic suppressor gene.

## Results

### LINC00844 is a novel androgen up-regulated prostate cancer associated lncRNA

To identify and characterize novel clinically relevant lncRNAs that are associated with prostate carcinogenesis, we processed and analyzed RNA sequencing (RNA-seq) expression data from two independent prostate cancer studies (Fig. 1a) [18, 19]. We first used DESeq2 [20] to obtain a list of differentially expressed transcripts (malignant vs benign or normal tissue) from each cohort and then used metaRNAseq [21] to perform meta-analysis on the two lists. The combined p-value from multiple testing was further corrected by the Benjamini and Hochberg method and using a false discovery rate (FDR) <0.01 we identified 370 clinically relevant lncRNAs (Table S1). Since our lab is interested in androgen-mediated transcription in prostate cancer cells, we decided to focus on the androgen-regulated lncRNAs within this group. To do this, we first defined a set of androgen-regulated lncRNAs with a minimum of 1.5-fold expression change from the RNA-seq data of LNCaP and VCaP cells that have been treated with or without dihydrotestosterone (DHT) [22]. Next, we overlapped this list of androgen-regulated lncRNAs with the differentially regulated list of lncRNAs from the prostate cancer studies. From this, we found 38 lncRNAs that are both clinically relevant and under androgen transcriptional regulation (Table S2). We further ranked this list based on their abundance in LNCaP and VCaP cells and validated the top 15 candidates by qPCR (Fig. 1b). For this study, we functionally characterized the lncRNA, LINC00844.

**Figure 1.**
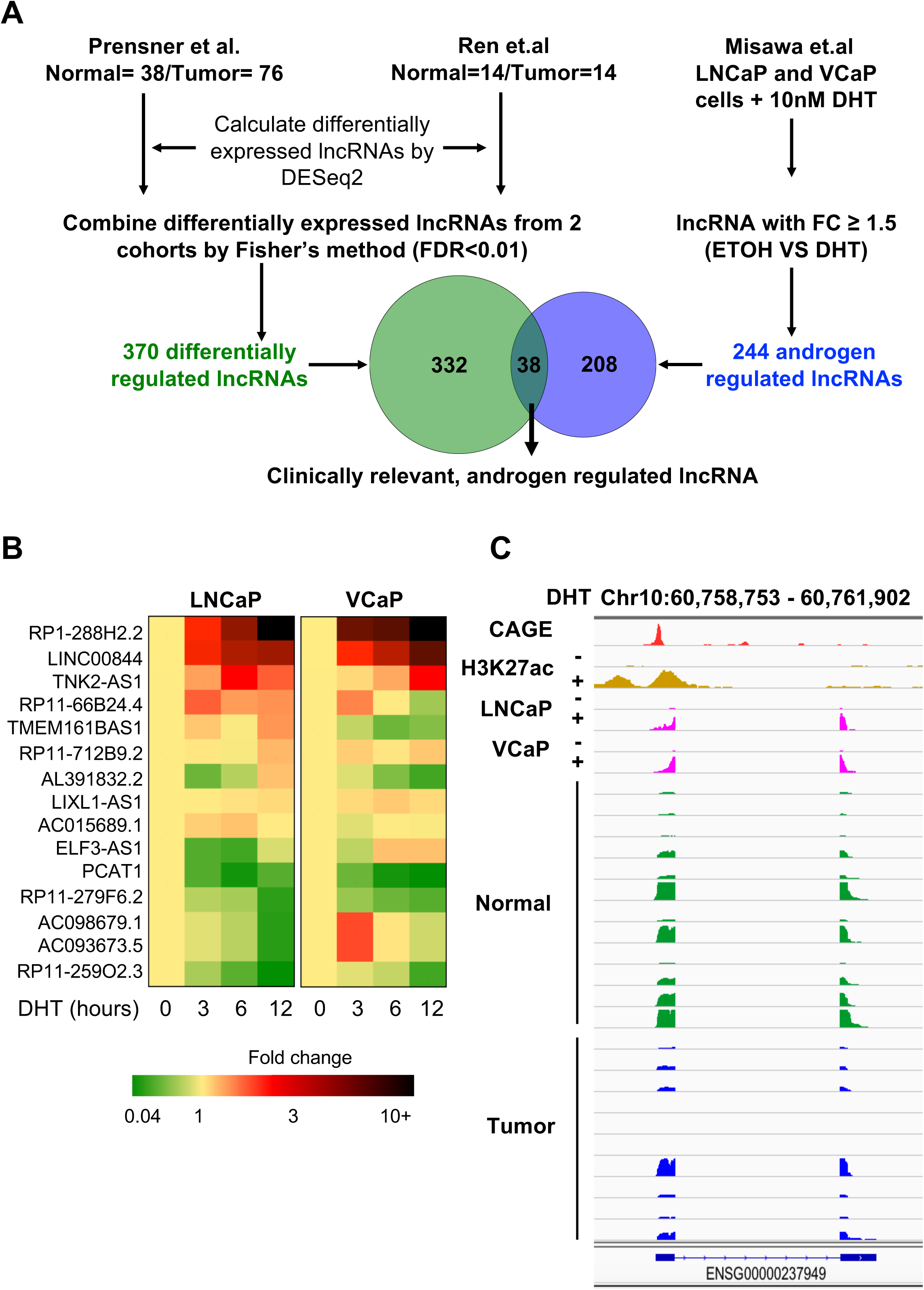
Identification and characterization of LINC00844 in prostate cancer cells. A) Workflow showing the steps in the discovery and identification of LINC00844. B) Heatmap representing the expression of 15 lncRNA in hormone starved LNCaP and VCaP cells stimulated with 10 nM of DHT for the indicated amount of time. C) Genome browser view of RNA-seq and ChIP-seq events around LINC00844 with the current annotation from Gencode. CAGE peaks from normal prostate tissues were obtained from the FANTOM database. H3K27ac ChIP-seq from LNCaP cells, RNA-seq from LNCaP and VCaP cells treated with ETOH or DHT were obtained from the GEO database (GSE51621 and GSE82223). RNA-seq data in normal and prostate cancer tissues were obtained from SRA database (ERP000550).

We began our characterization of LINC00844 by determining it’s coding potential to confirm whether it is indeed a non-coding transcript. We did this by examining the DNA sequence of LINC00844 using the Coding Potential Calculator (http://cpc.cbi.pku.edu.cn/) [23]. A score of −1.04818 was obtained for LINC00844 which suggests that open reading frame of this lncRNA lacks the potential to code for a protein. Similarly, when we examined the coding potential of HOTAIR, a well-known lncRNA, we obtained a score of −1.19011. In contrast, we obtained a score of 12.246 for GAPDH.

Next, we examined the basic features of the LINC00844 transcript. LINC00844 is an annotated lncRNA (ENSG00000237949, Entrez:100507008) transcribed from an intergenic region on Chromosome 10q21.1. The basic Genecode annotation maps it as a 477 bp transcript consisting of two exons transcribed from a region of 1.98 kb. To characterize LINC00844 further, we interrogated several publically available datasets including the recently published Cap Analysis of Gene Expression (CAGE) [24] on prostate tissues, H3K27 acetylation (H3K27ac) ChIP-seq in LNCaP cells [25], and the RNA-seq expression data from LNCaP and VCaP cells [26] as well as prostate cancer patient samples [19]. As shown in Fig. 1c, there is a strong H3K27ac peak at the TSS of LINC00844 (marked by the CAGE peak) which increases upon DHT stimulation. Moreover, there is a concomitant increase in the transcript level of LINC00844 in both LNCaP and VCaP cells. In patient samples, LINC00844 is expressed in normal and prostate cancer. Taken together, our results suggest that LINC00844 is a novel prostate cancer associated lncRNAs that is expressed in prostate tissues and is regulated by androgen in prostate cancer cells.

### LINC00844 is a direct AR target

Next, we assessed the expression and regulation of LINC00844 in different prostate cancer cell lines. LINC00844 transcript level is highest in the AR-dependent cell lines, LNCaP and VCaP, while it is minimally expressed in the AR sensitive cell line, 22Rv, and undetectable in the AR-independent cell line, PC3 (Fig. 2a). With respect to its regulation, LINC00844 was stimulated by DHT in a time- and concentration-dependent manner similar to KLK3/PSA, the model AR-regulated gene (Fig. 2b-c and Fig. S1a-b). In contrast, DHT did not significantly affect the expression of LINC00844 in 22Rv1 cells (Fig. S1c). To further validate the AR-mediated regulation of LINC00844, we treated LNCaP and VCaP cells prior to DHT stimulation with either anti-androgens (Bicalutamide or MDV-3100) or siRNA against AR. In both conditions, we found DHT-mediated stimulation of LINC00844 was greatly reduced (Fig. 2d–e).

**Figure 2.**
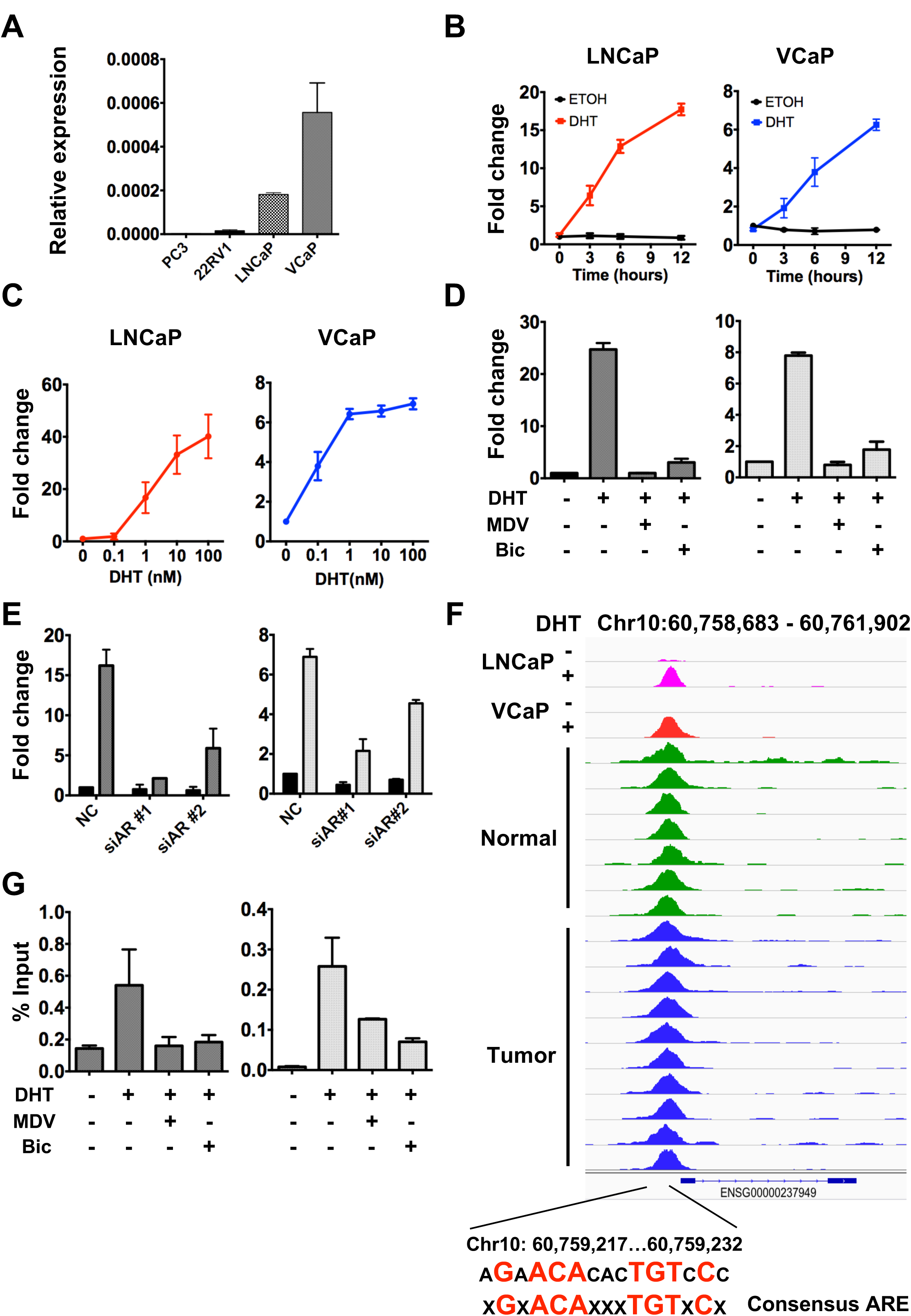
LINC00844 expression is directly regulated by AR. The relative expression of LINC00844 measured by qPCR in A) different prostate cancer cells maintained in normal media, LNCaP and VCaP cells treated with B) 10 nM of DHT for indicated time point, or C) different concentration of DHT for 12 h, or D) pre-treated with 10 µM of Biclutamide (Bic) or MDV-3100 (MDV) for 12 h followed by 10 nM DHT stimulation, or E) transfected with siAR followed by 10 nM DHT treatment for 12 h. F) Screenshot of AR ChIP-seq peaks mapped to the promoter of LINC00844 from LNCaP and VCaP cells treated with 100 nM of ETOH or DHT for 2 h and prostate cancer patient samples from GEO dataset (GSE70079). The bottom of the screenshot shows a zoomed in region of the AR binding site upstream of the LINC00844 TSS containing a canonical AR binding motif. G) AR ChIP-qPCR of the region spanning the LINC00844 promoter in LNCaP and VCaP cells pre-treated with 100 µM of Biclutamide (Bic) and MDV3100 (MDV) followed by 100 nM DHT stimulation.

To determine if AR directly regulates the expression of LINC00844 in prostate cancer by binding in the upstream or downstream regulatory region of the transcript, we interrogated AR ChIP-Seq datasets from cell lines [27, 28] and patient samples [29]. As shown in Fig. 2f, there is an ARBS located at the TSS of LINC00844 in LNCaP and VCaP cells as well as in patient samples. In addition, DHT enhanced the recruitment of AR to this site similar to other ARBS associated with known AR-regulated genes (Fig. S1d). We verified the binding of AR to the TSS of LINC00844 by ChIP-qPCR and showed that recruitment was abolished by antiandrogens in both LNCaP and VCaP cells (Fig. 2g). Upon examination of the DNA sequence of the ARBS, we identified the presence of a canonical AR response element (ARE) motif (Fig. 2f), suggesting that AR binds directly to this site. Taken together, our results suggest that LINC00844 is a direct AR-regulated lncRNA.

### LINC00844 is suppressed in malignant and metastatic prostate cancer

To begin understanding the clinical significance of LINC00844 in prostate cancer, we examined the expression profile of the transcript in patient samples. The expression level of LINC00844 is significantly lower in malignant tumor samples compared to their matched controls or benign samples in four prostate cancer cohorts [18, 19, 30], including the two cohorts that we had utilized for our initial characterization of lncRNAs (Fig. 3a-d). This result suggests a potentially important tumor suppressive functional role for LINC00844. Notably, two out of the four cohorts which also contain the expression information for metastatic prostate cancer samples showed the level of LINC00844 transcript in these metastatic samples are further suppressed (Fig, 3b and d). Taken together, our results show LINC00844 is the first AR-regulated lncRNA to demonstrate profound association with both malignant and metastatic prostate cancer.

**Figure 3.**
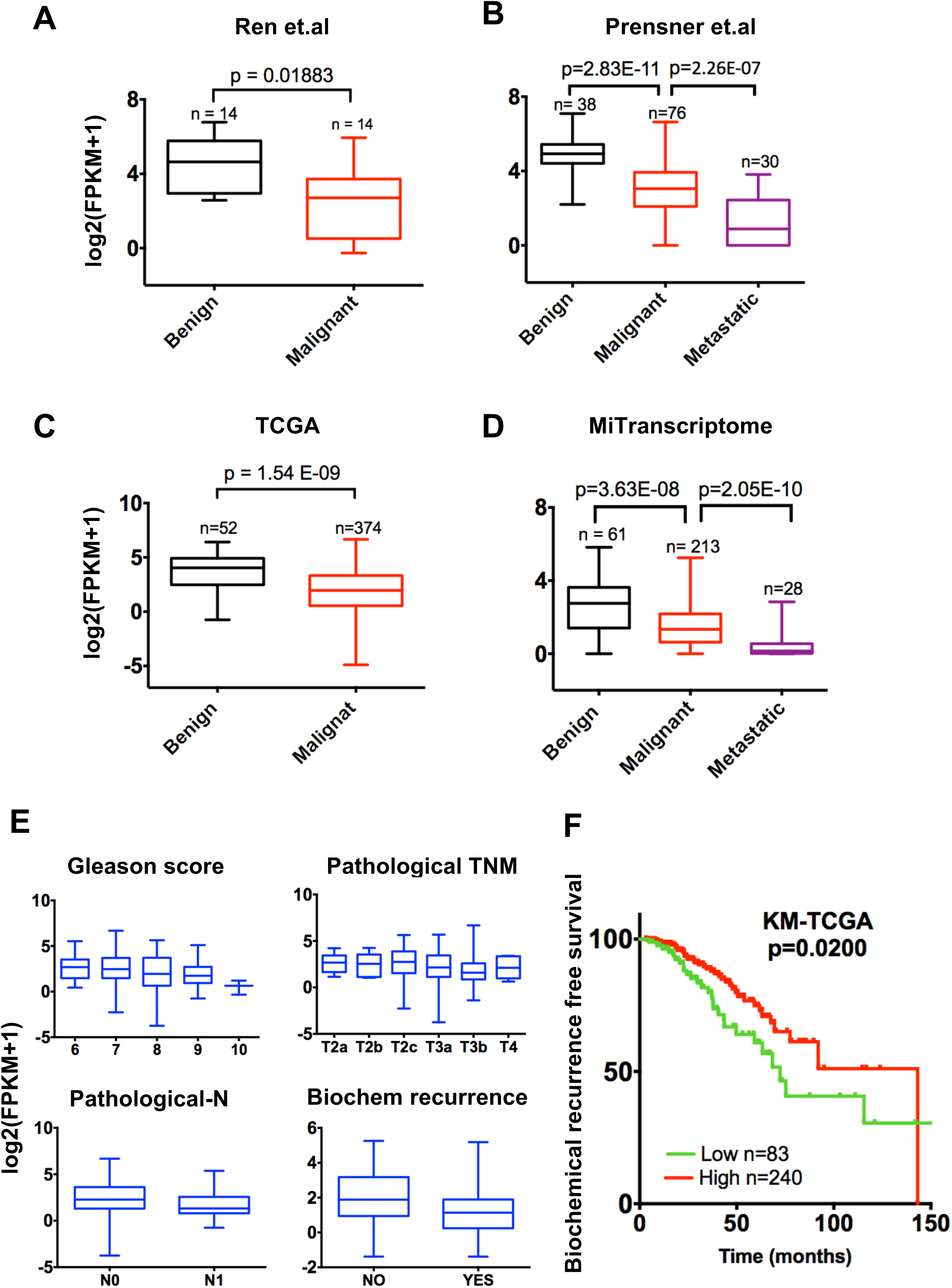
LINC00844 is suppressed in malignant and metastatic prostate cancer. Box-plot of LINC00844 expression presented as log_2_(FPKM) in published cancer cohorts, A) Ren et. al., B) Prensner et. al., C) TCGA-PRAD, and D) MiTranscriptome. E) LINC00844 expression in TCGA samples segregated based on clinical parameters including Gleason score, pathological TNM staging, spread of tumor to neighboring lymph node, and biochemical recurrence. F) Kaplan–Meier analysis of prostate cancer outcome. 384 prostate cancer samples from the TCGA dataset were separated as upper 3 quarters and lower 1 quarter based on the expression of LINC00844 and analyzed for period of biochemical recurrence free survival.

We further examined the association of LINC00844 expression level with additional clinical parameters such as Gleason score, spread of cancer to lymph nodes, and biochemical recurrence. As shown in Fig. 3e, there is a negative correlation between LINC00844 expression level and Gleason score (*i.e*. tumors with higher Gleason score which denotes more aggressive form of the cancer show the least expression for LINC00844). Similarly, tumors with lower expression of LINC00844 not only have higher rate of biochemical recurrence but also increased spread of the tumors to the neighboring or distant lymph nodes (Fig. 3e). Finally, we examined the expression level of LINC00844 with respect to prostate cancer patient recurrence free survival. As expected, patients with higher expression of LINC00844 had a significantly lower recurrence-free survival (log-rank p< 0.02) (Fig. 3f). Taken together, the clinical expression data of LINC00844 suggest it may not only be involved in prostate carcinogenesis but also in metastatic transformation.

### LINC00844 inhibits cell migration and invasion

Since the above clinical observations indicate that LINC00844 may play an important role in metastatic transformation of malignant prostate cancer, we decided to test this out by examining the effect of LINC00844 knock-down on LNCaP cell migration and invasion using the Boyden’s chamber assay. Knock-down of LINC00844 significantly increased both the migration (Fig. 4a, left panel) and invasion (Fig. 4b, left panel) of LNCaP cells cultured in normal growth media. Moreover, the increase in both invasion and migration was regulated by DHT (Fig. 4a and b, right panels), suggesting LINC00844 may mediate its effects through the AR signaling pathway. Next, we determined if the observed increase in cell migration and invasion was due to an increase in cell proliferation. As shown in Fig. 4c, we did not observe any significant increase in cell proliferation after LINC00844 depletion. While many reported lncRNAs in prostate cancer are known to affect cell proliferation and migration [26, 31], our results show LINC00844 affects exclusively prostate cancer cell migration and invasion, without affecting cell proliferation. Collectively, these results suggest that LINC00844 (i) is critical for preventing a metastatic phenotype in prostate cancer cells and (ii) mediates its effects through the AR signaling pathway.

**Figure 4.**
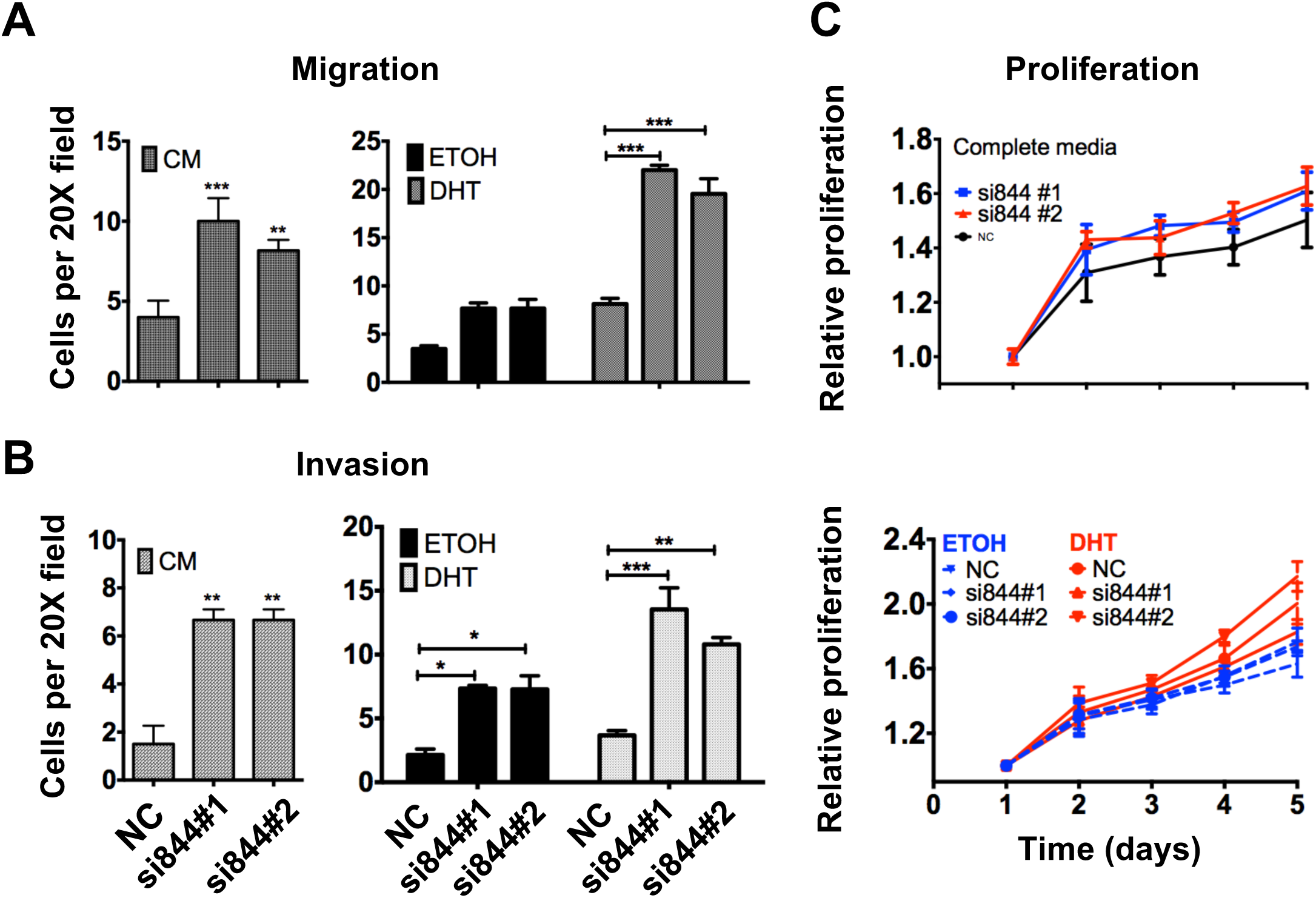
LINC00844 inhibits prostate cancer cell migration and invasion. Graph representing the number of LNCaP cells per high power field (20X objective) migrated across Transwell support with 8 µm pores. Transwell supports were coated with Matrigel in 1:20 dilution for B) invasion assay or left uncoated for A) migration assay. Error bars show SEM of cells counted in 5 high power fields of view in 3 independent experiments. C) Relative cell proliferation measured by Alamar blue assay in LNCaP cells transfected with siLINC00844 or NC control. Normalized Alamar blue emission at 560EX nm/590EM nm measured in triplicates from 3 independent experiments. Data represented as mean ± SEM. (*p<0.05, **p<0.01, ***p<0.001)

### LINC00844 is an integral component of the AR transcriptional network

To explore whether LINC00844 has a role in regulating AR signaling in prostate cancer, we performed microarray analysis on siNC or siLINC00844 treated LNCaP cells before and after DHT stimulation. Overall, we detected 917 DHT-responsive genes (fold change ≥ 1.5 and p <0.01) with 529 up- and 388 downregulated genes (Fig. 5a). Among the 917 DHT-regulated genes, 520 genes (327 up and 193 down) were affected by LINC00844 knock-down (fold change ≥ 1.2). With respect to the DHT-upregulated genes that were affected, 316 genes were repressed, while 11 genes were increased. Moreover, this group of DHT-upregulated genes included well-known model genes such as KLK2, KLK3, FKBP5, and GREB1. As for the DHT-downregulated genes, 177 were upregulated, while 16 were further suppressed. We validated our microarray results in both LNCaP and VCaP cells (Fig. 5c and d). In general, the depletion of LINC00844 suppressed the activation of androgen-regulated genes in both cell lines although the effects were more profound in LNCaP cells. Thus, our findings suggest that LINC00844 is important regulator of androgen-dependent gene transcription.

**Figure 5.**
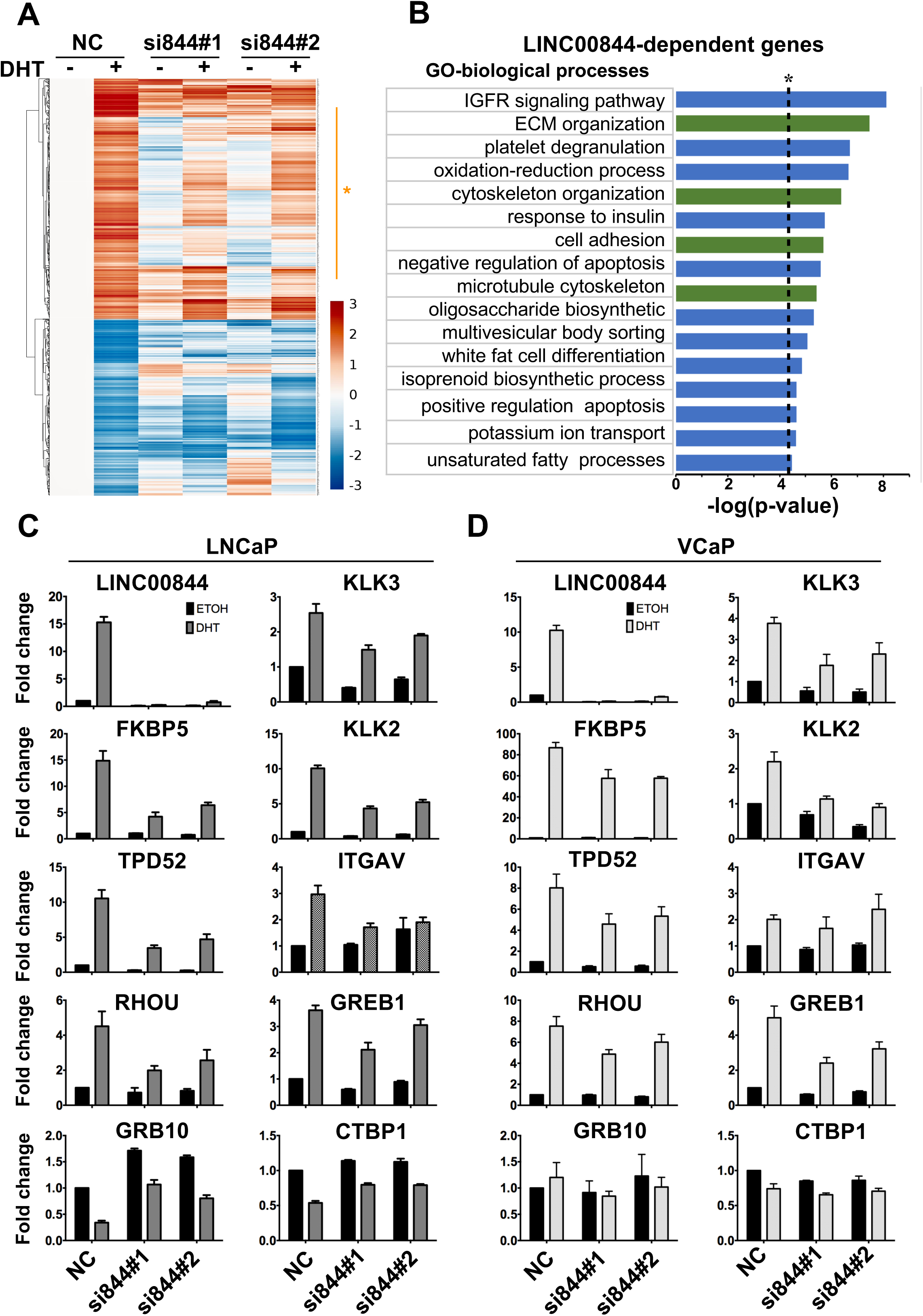
LINC00844 regulates a subset of AR target genes. A) Gene expression profiling was performed in LNCaP cells transfected with NC or siLINC00844 with DHT or ETOH stimulation for 12 h. Heatmap represents all the AR-regulated genes (NC-ETOH vs NC-DHT ≥ 1.5 p<0.01) and their corresponding expression in siLINC00844 condition. The fold change in expression is mentioned below. The AR-upregulated genes that were suppressed by ≥ 1.2-fold after LINC00844 knock-down are represented by orange asterisks. B) GO terms associated with biological processes enriched in LINC00844-dependent genes. The dotted line represents p<0.05. The GO terms associated with cell migration and invasion are highlighted in green. C and D) qPCR validation of several selected genes form the microarray list identified to be regulated by LINC00844. Results are shown as the mean fold change + SEM from at least 3 independent experiments.

We next performed GO analysis using DAVID Functional Annotation Bioinformatics Microarray Analysis (https://david.ncifcrf.gov) [32] on the above androgen-regulated genes that we identified as dependent on LINC00844. In support of our functional studies, we found that androgen-regulated genes that are dependent on LINC00844 for expression are enriched in processes associated with cytoskeletal organization, extracellular matrix organization, microtubule cytoskeleton organization, cell adhesion, structural component of cytoskeleton, and anatomical structural morphogenesis, all of which are GO terms linked to cell migration or metastasis (Fig. 5b). Taken together, our results indicate that LINC00844 is an integral component of the AR transcriptional network and a decrease or loss of LINC00844 expression may promote malignant transformation of prostate cancer by suppressing the DHT-mediated activation of genes.

### LINC00844 regulates AR binding to chromatin

Next, we focused on understanding the underlying mechanism on how LINC00844 might control the expression of androgen-regulated genes. Previously, our lab and others have demonstrated that AR collaborative factors such as NKX3-1 [27] and lncRNAs like PlncRNA [33, 34] act through a feed forward mechanism to regulate AR mRNA and protein levels to influence the expression of downstream AR target genes. Thus, we examined if LINC00844 could also function in a similar fashion. As shown in Fig. 6a and b, knock-down of LINC00844 did not alter the mRNA or protein level of AR in both LNCaP and VCaP cells, indicating that LINC00844 likely functions by directly or indirectly influencing the activity of AR, such as through chromatin binding.

**Figure 6.**
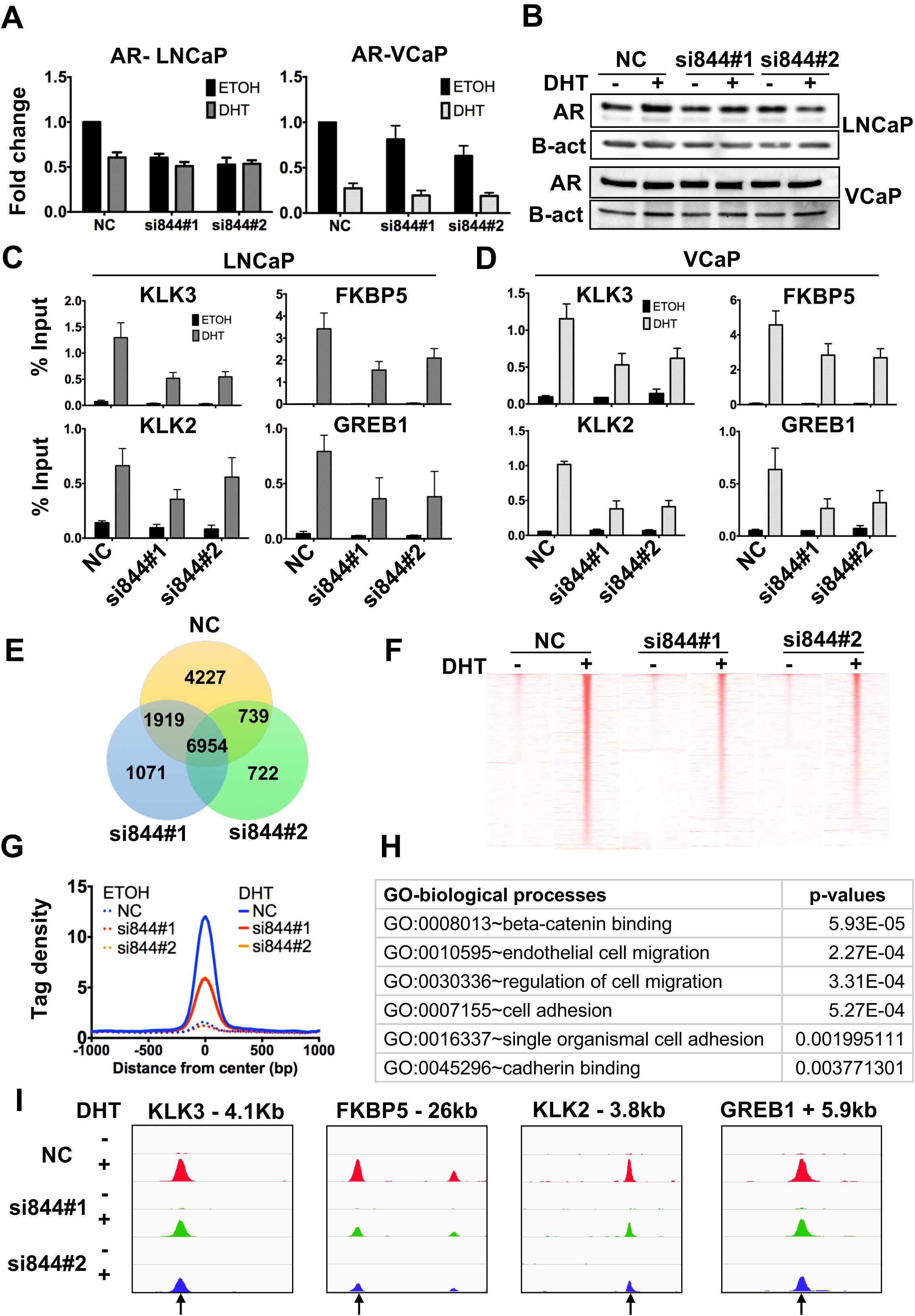
LINC00844 promotes AR association to the chromatin. A-B) AR mRNA and protein levels were quantified by qPCR and western blot in LNCaP and VCaP cells treated with siLINC00844 or NC and stimulated with 100 nM of DHT for 2 h. C-D) AR ChIP-qPCR was performed in LNCaP and VCaP cells transfected with siLINC00844 or NC and treated with 100 nM of DHT for 2 h, graph represents AR enrichment at the ARBS within +/− 50 kb from the TSS of the specified gene. All results represent the average of 3 individual experiments ± SEM. E) Venn diagram representing the common ARBS identified from 2 independent experiments in LNCaP cells treated with siLINC00844 or NC and 100 nM of DHT for 2 h. F) A heatmap representing the sorted AR ChIP-seq signal corresponding to NC-DHT specific regions in all the tested conditions. Signals are plotted in reference to the center of AR ChIP-seq cluster peak (-/+ 1 kb). G) A graph comparing the average ChIP-seq tag density around ± 1 kb from the AR peak center in NC-DHT compared to siLINC00844-DHT (solid lines) and NC-ETOH compared to siLINC00844-ETOH (dotted lines). H) A table listing the GO terms associated with cell migration that are enriched in the ARBS specific to NC-DHT was calculated by GREAT tool. I) Snapshots showing the AR ChIP-seq peak at the enhancer region of 4 model AR-regulated genes (KLK3, FKBP5, KLK2, and GREB1). Black arrow represents the ARBS with marked reduction in AR binding after LINC00844 knock-down.

Hence, we next determined whether LINC00844 is important for the recruitment of AR to ARBS by performing chromatin immunoprecipitation (ChIP) assays. For this, we began by examining the ARBS associated with KLK2, KLK3, FKBP5, and GREB1, AR-regulated genes whose expression we have shown is dependent on LINC00844. We tested the recruitment of AR to these binding sites in LNCaP and VCaP cells that were treated with siLINC00844 or siNC. Interestingly, AR recruitment at these regions was markedly reduced by LINC00844 depletion (Fig. 6c and d). These results suggest that LINC00844 influences the expression of the AR-regulated transcriptome in part by facilitating the recruitment of AR to the chromatin.

To examine whether LINC00844 has a global effect on AR binding, we performed AR ChIP-seq on LNCaP cells that have been treated with or without siLINC00844. The experiment was performed in duplicates for increased confidence and we considered only the common ARBS for down-stream analysis. Overall, the depletion of LINC00844 resulted in a genome-wide reduction of a large number of ARBS (Fig. 6e and Table S3). Specifically, 4,227 ARBS (~30%) in the NC-DHT condition were lost after LINC00844 knock-down. Moreover, the ChIP-seq tag intensity of these 4,227 ARBS were markedly reduced after LINC00844 depletion (Fig. 6f, g, and i). GREAT analysis of the ARBS that were affected by LINC00844 depletion showed many terms enriched for cell migration which is consistent with our initial finding that LINC00844 regulate the expression of genes associated with prostate cancer cell migration and invasion (Fig. 6h). Taken together, our results suggest that LINC00844 regulates global androgen-dependent transcription in *trans* by modulating the binding of AR to chromatin.

### LINC00844 inhibits prostate cancer cell migration and invasion by upregulating the expression of NDRG1

Next, to identify the potential target gene(s) responsible for mediating the LINC00844 associated phenotype, we performed in-depth analysis of the androgen-regulated genes that requires LINC008844 for expression. We specifically concentrated on the genes constituting the GO terms enriched for migration and invasion processes. From this analysis, we identified and validated N-Myc Downstream Regulated 1 (NDRG1) as a DHT- and LINC00844-dependent gene (Fig. 7a). In addition, we showed that knock-down of LINC00844 reduced the binding of AR to an ARBS located ~30 kb upstream from the TSS of NDRG1 (Fig. 7b and c).

**Figure 7.**
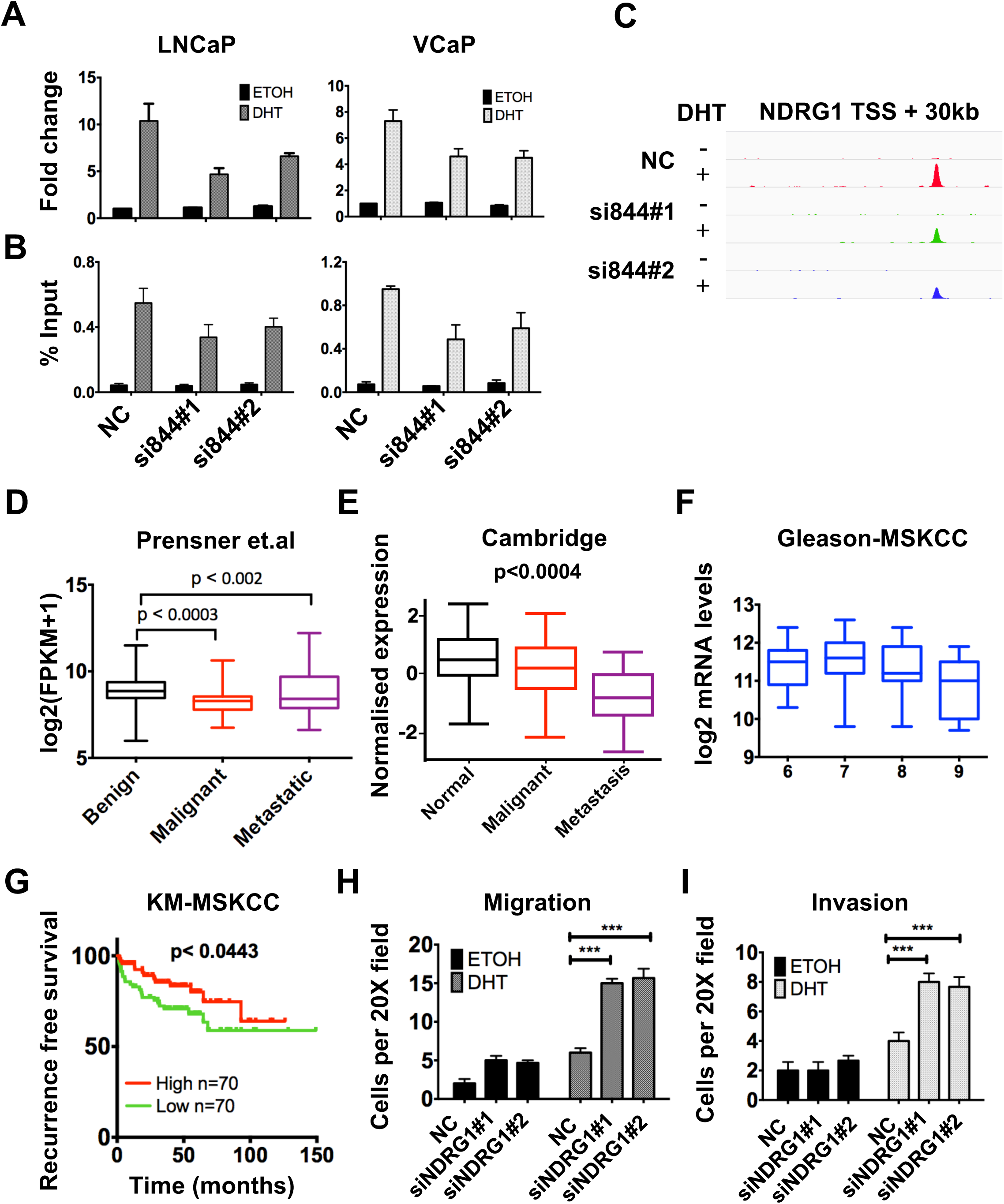
LINC00844 mediates the phenotypic effects in part by regulating NDRG1 expression. A) Bar graph showing the expression of NDRG1 measured by qPCR in LNCaP and VCaP cells treated with siLINC00844 or NC and 10 nM of DHT or ETOH for 12 h. B) AR ChIP-qPCR was performed in LNCaP and VCaP cells treated with siLINC00844 or NC and 100 nM of DHT or ETOH for 2 h. Graph representing AR binding to the ARBS at the enhancer region of NDRG1 gene (30 kb form the TSS) and C) a screenshot showing the AR ChIP-seq peak from the same region in LNCaP cells. Results are shown as the mean fold change ±SEM from at least 3 independent experiments. Box-plot showing the expression of NDRG1 in D) Prensner et.al as log2(FPKM+1), E) Cambridge study dataset as average expression, and F) patients in MSKCC datasets segregated based on their Gleason score and presented as log2mRNA expression values. G) Kaplan–Meier analysis of prostate cancer outcome. 140 patients from the MSKCC dataset were divided as higher or lower than median NDRG1 expression and analyzed for period of biochemical recurrence free survival. Graph representing the number of LNCaP cells per high power field (20X objective) migrated across Transwell supports with 8 µm pores. Transwell supports were coated with Matrigel in 1:20 dilution for I) invasion assay or left uncoated for H) migration assay. Error bars show SEM of cells counted in 5 high power fields of view in 3 independent experiments. (*p<0.05, **p<0.01, ***p<0.001)

NDRG1 is associated with a number of different types of cancers including breast, colon, pancreas, and prostate cancer [35–39]. In addition, NDRG1 expression is negatively correlated with the cancer status and overexpression studies with NDRG1 in prostate cancer cells markedly reduced *in vivo* metastasis [38]. NDRG1 expression is significantly reduced in prostate tumors with lymph node or bone metastasis compared with localized prostate cancer [38, 40, 41]. Interrogation of data from the study by Prensner et.al [18] and a recently published work from the CamCaP group [42] showed a reduction in NDRG1 expression in malignant and metastatic samples as compared to normal tissue (Fig. 7d and e). Similarly, NDRG1 expression was also negatively correlated with Gleason score (Fig. 7f) and low expressing patients showed significantly low recurrence-free survival in MSKCC dataset [43] (Fig. 7g).

From the above results, we hypothesized that LINC00844 may control the invasiveness of prostate cancer cells in part by regulating the expression of NDRG1. To test this possibility, we transfected LNCaP cells with siNDRG1 and assayed for their migration and invasion ability. If the activation of NDRG1 is essential for inhibiting cell migration and invasion, then we should observe an increase in migration and invasion upon the knock-down of NDRG1. Indeed, depletion of NDRG1 significantly increased both the migration (Fig. 7h) and invasion (Fig. 7i) activity of LNCaP cells. Collectively, our results suggest that AR and LINC00844 control prostate cancer metastasis by directly regulating the transcription of NDRG1.

## Discussion

Recent discoveries and advancements in transcriptomics have revealed crucial roles for the so called “junk DNA” or ncRNA in a myriad of biological processes including cancer [11]. To further our understanding of ncRNAs in prostate cancer, we have systematically analyzed two well established prostate cancer cohorts of different ethnicity to identify and characterize a novel lncRNA, LINC00844.

AR is critical for the normal development of prostate tissue and plays an important role in its carcinogenesis [3]. AR-mediated transcriptional regulation is central in defining the transcriptomic landscape associated with different stages of prostate cancer like hormone sensitive and metastatic CRPC. The AR regulatory complex consists of various pioneer factors, co-activators, and co-repressors that are recruited to gene promoters in a coordinated and sequential pattern [4]. Recent discoveries have exemplified the critical role of lncRNAs in this complex, such as mediating AR long-range chromatin interactions and altering the chromatin confirmation to regulate AR binding to the chromatin [15]. Yet the current list of lncRNAs associated with AR transcriptional complex still remains small. In this study, we demonstrate LINC00844 is a novel AR-regulated and prostate cancer-associated lncRNA that is required for regulating the core AR transcriptional network.

Currently, a limited number of prostate cancer associated lncRNAs have been identified and only a fraction of them have been studied in detail to divulge the underlying molecular mechanisms. Interestingly, most of the lncRNAs studied so far are known to be upregulated in malignancy, however, our findings show LINC00844 is suppressed in malignancy which suggests a tumor suppressive functional role. In this work, we showed LINC00844 is significantly suppressed not only in the two test cohorts [18, 19] but also in much larger validation datasets, including TCGA [44] and MiTranscriptome [30]. Interestingly, we also found the expression of LINC00844 is further suppressed in metastatic prostate cancer samples which suggest that it be may be crucial for inhibiting the metastatic transformation of malignant prostate cancer. In support of a tumor suppressive functional role for LINC00844, we observed that prostate tumors with low expression of LINC00844 have worst prognostic outcomes and high biochemical recurrence.

Similarly, by means of loss-of-function studies we demonstrated that the loss of LINC00844 resulted in increased cell migration and invasion without affecting the rate of proliferation. Interestingly, we also observed that the effect of LINC00844 was more pronounced on DHT-mediated cell migration and invasion compared to cells grown in normal growing media or ETOH stimulated cells. Hence, we argue that LINC00844 mediates its phenotypic effects on prostate cancer cells by primarily targeting the AR transcriptional network.

Our current work indicates that LINC00844 is a critical component of the AR transcriptional machinery. From our microarray analysis, we showed that a loss of LINC00844 leads to a suppression of a subset of DHT-activated genes that are mainly associated with cytoskeletal organization or proliferation of prostate processes. Mechanistically, LINC00844 appears to be crucial for the binding of AR to the chromatin. To probe deeper into the mechanism through which LINC00844 regulates the binding of AR to the chromatin, we performed RNA immunoprecipitation (RIP) assays to examine if LINC00844 physically interacts with AR. Surprisingly, we did not find evidence of any direct physical interactions between LINC00844 and AR (data not shown). Hence, at this stage we can only speculate an indirect regulation of AR by LINC00844. Plausible candidates could be coactivators or collaborative factors of the AR, however, this will need to be further studied.

We also provided evidence to show that LINC00844 could potentially regulate migration and invasion of prostate cancer through NDRG1. A previous study has shown that a loss of NDRG1 expression significantly increases *in-vivo* metastasis of prostate cancer cells [38]. Similar to LINC00844, NDRG1 is also significantly suppressed in malignant and metastatic prostate cancer samples and patients with low expression have significantly increased rates of recurrence. Herein, we demonstrated *in vitro* DHT-mediated activation of NDRG1 suppresses prostate cancer cell migration and invasion and LINC00844 is critical for AR recruitment to its enhancer and its expression.

In summary, our work showed LINC00844 is a novel AR-regulated lncRNA that plays an important role in the regulating prostate cancer cell migration and invasion. In addition, LINC00844 regulates AR recruitment to chromatin and enhances the activation of a large number of canonical target genes.

## Acknowledgement

This work was supported by a University of Macau Multi-Year Research Grant (MYRG2015-00196-FHS), a University of Macau Start-up Research Grant (SRG2014-0000-FHS), and the Macau Science and Technology Development Fund (FDCT/023/2014/A1 and FDCT102/2015/A3) to EC.

## Material and Methods

### Reagents and antibodies

Dihydrotestosterone (DHT) was purchased from Tokyo Chemical Industry, MDV-3100 and Biclutamimide were purchased form TCI Chemicals, USA. The following antibodies were used for ChIP and western blot analyses: anti-AR (sc-815X, sc-H280 and sc-816).

### Cell culture

LNCaP, VCaP, and 22Rv1 were purchased from ATCC. LNCaP and 22Rv1 cells were maintained in RPMI medium 1640 (RPMI) (Gibco) supplemented with 10% fetal bovine serum (FBS) (Gibco). VCaP cells were grown in Dulbecco’s modified Eagle medium (DMEM) (Gibco) supplemented with 10% FBS, 1 mM sodium pyruvate, 0.08% sodium bicarbonate. Prior to Ethanol (ETOH) or DHT treatment, LNCaP and 22Rv1 cells were deprived of hormones for 3 days by growing them in phenol red-free RPMI containing 10% charcoal-dextran-treated FBS (HyClone), whereas VCaP cells were grown for at least a day in phenol red-free DMEM containing 10% charcoal-dextran treated FBS.

### siRNA studies

LNCaP cells cultured for 24 h in phenol red-free medium were transfected with 5 nM of Dicer-substrate short interfering RNAs (siRNAs) from IDT, using Lipofectamine RNAiMAX transfection reagent (Invitrogen) according to the manufacturer’s protocol. After 24 h of incubation, the cells were transfected again in a similar manner with 5 nM siRNA. 48 h after the second round of transfection, cells were treated with ETOH or 10 nM DHT for another 12 h before harvesting for real-time reverse transcription-qPCR (RT-qPCR), microarray, and western blot analyses. For every target, 2 separate siRNA were paired with the control nontargeting siRNA (NC). siRNA and real-time qPCR primer sequences are listed in Table S4 in the supplemental material. Gene expression profiles for knockdown studies were obtained from at least three independent experiments.

### Chromatin immunoprecipitation (ChIP)-qPCR and ChIP-seq

ChIP assays were performed with minor modifications to the protocol described by Carroll and colleagues [45]. For knock-down ChIP assays, 3 million LNCaP cells or 5 million VCaP cells were transfected twice with siRNA targeting LINC00844 or control. 48 h after transfection and 2 h of 100 nM DHT/ETOH treatment, cells were crosslinked with 1% formaldehyde and harvested. Cells were lysed, nuclei extracted and DNA sonicated using a Biorupter (Diagenode) before overnight immunoprecipitation with AR antibody conjugated to protein G magnetic beads (Dynabeads, Invitrogen). The beads were thoroughly washed and reverse crosslinked at 65^o^C along with 5% input before isolating DNA using a PCR-purification kit (Qaigen). AR enrichment was quantified by real-time qPCR using primer sequences listed in Table S3 and represented as percentage of input.

For the high throughput sequencing of the ChIP DNA (ChIP-seq), DNA libraries were constructed using the NEBNext^®^ Ultra™ DNA Library Prep Kit for Illumina^®^ (NEB, catalogue # E7370L) according to manufacturer’s protocol. The libraries were amplified for 12 cycles by PCR and purified using AMPure XP beads (Beckman Coulter, catalogue # A63880). The barcoded libraries were mixed in equimolar ratio and sequenced using the Illumina HiSeq-2000. Sequencing reads were mapped with BWA [46] to the indexed reference genome UCSC hg19. The obtained BAM files (as replicates) were subjected for peak calling using MAC2 [47] using default parameters. Specific and overlapping peaks between different test conditions were identified using the HOMER package [48]. Heatmaps of the read density and the average binding profiles around the center of peaks (±1 kb) were produced using the R heatmap package. Genome coverage bedgraph files were converted to bigwig format using the bedGraphToBigWig tool and loaded onto the IGV browser to obtain screenshots. The raw sequencing data can be found at the GEO with the accession number GSE108704.

### Migration and invasion assay

Migration and invasion assays were carried out using LNCaP cells transfected with siRNA and seeded in Boyden’s Transwell Chamber inserts (uncoated for migration and 1:20 diluted Matrigel coating for invasion) in RPMI media containing 5 nM of siRNA in 100 µl of RPMI with 1% FBS. 600 µl of RPMI with 10% FBS containing ETOH or DHT in the lower chamber acted as chemoattractant. After 48 h, cells were fixed with 4% formalin and stained with crystal violet and cells migrated to the other side of the insert were counted using an inverted bright field microscope at 20X magnification.

### Alamar blue cell proliferation assay

LNCaP cells transfected with siRNA for 24 h were seeded in multiple 96 well plates (as triplicates) at a concentration of 10,000 cells/well followed by a second round of siRNA transfection. To assess for proliferation, cells were incubated with ALAMAR blue reagent for 3 h and the fluorescence was measured by the Spectramax with 560EX nm/590EM nm filter settings. The results from 3 independent experiments are presented as mean ± S.E.M

### Microarray expression profiling

Purified total RNA from three independent biological replicates were converted to cRNA using the TargetAmp™-Nano Labeling Kit for Illumina Expression BeadChip (Epibio) according to the manufacturer’s instructions. cRNA was hybridized onto HumanHT-12 v4 Expression BeadChips (Illumina). The BeadChips were scanned with the iScanner (Illumina) and the image data was processed using GenomeStudio. The gene expression data was analyzed using Lumi based tools from www.arrayanalysis.org [49]. All the microarray data has been deposited at the GEO database with the accession number, GSE*** (will be provided later).

